# YggG is a Novel SPI-1 Effector Essential for *Salmonella* Virulence

**DOI:** 10.1101/300152

**Authors:** Menghan Li, Bing Gu, Rajdeep Bomjan, Meghana Chitale, Daisuke Kihara, Daoguo Zhou

## Abstract

*Salmonella* is a leading cause of food borne illness and poses a major public health problem worldwide. Two type III secretion systems (TTSSs) are responsible for the delivery of a series of bacterial effectors into the infected cells to reprogram host cell functions to promote bacterial invasion and intracellular survival/replication. Nearly half of *Salmonella* genes encode proteins that are annotated as “hypothetical”. We hypothesize that some of these hypothetical proteins might be TTSS effectors and participate in *Salmonella* virulence. In this study, we employed an *in silico* screen to identify putative TTSS effector proteins and performed a large scale screening of *Salmonella* effectors using a β-lactamase protein translocation reporter assay. We identified 22 novel effectors that have not been previously reported. One of the effector – YggG, is found to be cytotoxic to mammalian cells when overexpressed suggesting interference with mammalian cell functions and survival. Importantly, *Salmonella* strains lacking YggG has reduced virulence in the mouse infection model. Our study demonstrated that YggG and its protease activity are required for *Salmonella* virulence in the mouse infection model. Surprisingly, no detectable roles in invasion into epithelial cells and no effect on the survival and replication inside cultured macrophages. We speculate that YggG may be involved in altering some aspects of the host immune system to promote infection. Our finding significantly expands the number of known *Salmonella* effectors, and laid a solid foundation in further understanding *Salmonella* pathogenesis.

**Abstract Importance:** Salmonellosis continues to be public health concern in the USA and the world. Salmonella delivers a set of bacterial proteins (effectors) into the host cells to invade non-phagocytic cells and to replicate inside the infected cells. Identification and characterization of these effectors are pivotal to the understanding of how *Salmonella* causes diseases. In addition, Nearly half of *Salmonella* genes encode proteins that are annotated as “hypothetical”. We have identified 22 novel effectors that have not been previously reported. Our finding significantly expands the number of known *Salmonella* effectors, and laid a solid foundation in further understanding *Salmonella* pathogenesis.

## Introduction

*Salmonella enterica* Serovar Typhimurium (*S*. Typhimurium) is a Gram negative, facultative intracellular bacterium that causes food borne illness and poses a major public health problem worldwide. *S*. Typhimurium possesses an arsenal of genes that are essential to establish a successful infection within the host. Many of the virulence traits of *S*. Typhimurium are directly linked to genes encoded within large regions of the bacterial chromosome titled *Salmonella* pathogenicity islands (SPIs). Two type III secretion systems (TTSS) encoded in SPI-1 and SPI-2 are responsible for the delivery of a panel of bacterial effectors into the host cells to reprogram host cell functions to facilitate bacterial invasion and survival/replication respectively (1).

*S*. Typhimurium invade host cells by inducing dramatic actin cytoskeleton rearrangements and membrane ruffling prior to bacterial entry (2, 3). SPI-1 plays an essential role in promoting *S*. Typhimurium invasion. SPI-1 secreted *<u>S</u>almonella* <u>i</u>nvasion <u>p</u>roteins (Sip) B-D are inserted into the host cell membrane to form the translocon, which are responsible for the transfer across host cell membrane of additional effectors into the host cells (4). These effectors, such as SipA, SipC, *<u>S</u>almonella* <u>o</u>uter <u>p</u>rotein (Sop)B, SopE and SopE2, function to induce the actin cytoskeleton rearrangement (membrane ruffling) leading to bacterial uptake. The biochemical and cellular mechanisms utilized by these effectors have been intense research subjects. SipA and SipC bind and modulate actin dynamics directly to promote actin polymerization and bacterial invasion (5–7). SopE and SopE2 modulate cytoskeleton rearrangement through binding to CDC42 and Rac1 and act as potent nucleotide exchange factors for these small G proteins, thereby activating CDC42 and Rac1 signal transduction pathways (8, 9). SopB has dual 4- and 5-phosphatase activities to regulate phosphoinositide dynamics to promote bacterial entry (10–14).

Upon entry into host cells, *Salmonella* resides in a vacuolar compartment termed the *<u>S</u>almonella* <u>c</u>ontaining <u>v</u>acuole (SCV). The SCV matures through a process whereby it selectively acquires endosomal components through vesicular fusion events and traffics towards the perinuclear region (15). Within the SCV, *Salmonella* is able to avoid the oxidative killing by the macrophages, thus facilitates the systemic spread of the infection (16). SPI-2 effectors have been shown to be responsible for SCV formation and maintenance, as well as the survival within the host. The precise mechanism of how SPI-2 effectors function to achieve this is not well understood.

Many of host cell functions modulated during *Salmonella* infection are dependent on the TTSS. A number of effectors have been characterized to be responsible for promoting bacterial invasion and survival/replication following entry as described above (3, 10–12, 16–20). Despite these advancements, factors responsible for many TTSS-dependent host responses, such as delayed macrophage cytotoxicity (21), avoidance of oxidative burst (17, 22–24), and altered inducible nitric oxide synthase (iNOS) localization (25–27), have yet to be defined. Therefore, elucidation of additional effectors may reveal the mechanisms governing these responses (17). For example, Hernandez *et al.* (2004) stated that *Salmonella* devoid of all known SPI-1 TTSS effector proteins retain their ability to induce cytotoxicity in caspase-1-deficient macrophages, while *Salmonella* lacking the InvG protein (a component of the SPI-1 encoded TTSS apparatus; essential for *Salmonella* SPI-1 effector protein secretion) completely lose this activity (28). This suggests that more effectors may exist mediating these responses. Furthermore, hundreds of *Salmonella* genes are annotated to encode “hypothetical” proteins. We speculate that some of the hypothetical proteins may be virulence factors and play important roles in *Salmonella* host cell interactions.

The structural components that made up the type III secretion apparatus share sequence similarities and can usually be identified by bioinformatics analysis. In contrast, the effectors that are secreted have no apparent sequence similarity, making them hard to identify (29–31). In an effort to identify additional TTSS effectors, we searched the *Salmonella* genome for proteins that matched the Gene Ontology (GO) terms that are likely to be required during *Salmonella* infections. A total of 383 putative effectors were identified and each was experimentally tested using a translocation reporter system. These efforts led to the identification of 22 novel TTSS-dependent effector proteins. One of the new effector, YggG, was found to be required for *Salmonella* virulence in a mouse infection model.

## Materials and Methods

### Bacterial strains and mammalian cell lines

*Salmonella* Typhimurium strains used in this study are isogenic derivatives of wild-type (WT) strain SL1344 (32). In-frame chromosomal gene deletions of *Salmonella* were generated using an allelic-exchange suicide vector pSB890 (33). Briefly, DNA fragment with the in-frame deletion was cloned into the suicide vector pSB890. Plasmids were introduced into *Salmonella* by conjugation, where they were integrated into the chromosome through homologous recombination. Each putative effector gene was fused in-frame with the 5’ of the TEM1 gene using the gateway clone strategy (Invitrogen, NY). Point mutation was generated using a QuikChange mutagenesis kit (Agilent Technologies, DE). DNA oligomer primers for PCR reactions are listed in **Table S1**.

*Escherichia coli* and *Salmonella* strains were routinely cultured in Luria-Bertani (LB) broth. *Salmonella* strains were cultured under SPI-1 TTSS-inducing conditions (LB broth with 0.3 M NaCl) for all of the invasion experiments. When applicable, antibiotics were used at the following concentrations: ampicillin 120 μg ml^−1^, streptomycin 25 μg ml^−1^, kanamycin 40 μg ml^−1^, and tetracycline 12 μg ml^−1^. Mammalian cell lines HeLa (CCL-2) and J774 were purchased from the ATCC cell biology stock center (Manassas, VA). HeLa and J774 cells were maintained in Dulbecco’s modified Eagle’s medium supplemented with 10% fetal bovine serum.

### β-lactamase assay TEM translocation assay

*Salmonella* strains expressing the putative effector-TEM1 fusions were used to infect monolayers of HeLa cells seeded in 96-well plates at an MOI of 20. 15min after infection, CCF4-AM, the substrate for beta-lactamase (Invitrogen, Carlsbad, CA) was added into the infected cells. Upon cleavage of CCF4, blue fluorescence will be observed. After further incubation for 2 hours at room temperature, infected cells were visually inspected under fluorescence microscope. The percentage of blue emitting cells was evaluated to identify translocation-positive clones. Experiments were performed in triplicate and at least 300 cells were counted for every infection.

### CyaA protein translocation assay

Plasmids expressing the CyaA fusion proteins were introduced into *Salmonella* SL1344. *Salmonella* expressing these fusions were used to infect HeLa cells at an MOI of 20 for 1 hour. Infected cells were washed 3 times with PBS, lysed with 0.1M HCl for 20min and centrifuged at 600x g for 5 min. Supernatant was taken for cAMP measurement using the cAMP Complete ELISA kit from Enzo Life Sciences (Farmingdale, NY.) according to the manufacturer’s instructions. Data presented are means ±SD from three independent experiments.

### Bacteria Invasion and replication Assays

*Salmonella* infections of HeLa cells were carried out as previously described (34). Briefly, *Salmonella* were cultured to an optical density of 1.0 (600 nm) in LB broth supplemented with 0.3 M NaCl at 37°C. Bacteria were then added to HeLa cells at an MOI of 10 and incubated for 15 min at 37°C in 5% CO_2_. After a defined period of incubation, infected cells were washed twice with phosphate-buffered saline (PBS) to remove extracellular bacteria and incubated further in DMEM containing 10% fetal bovine serum and 16 μg of gentamicin ml^−1^. At different time points after gentamicin treatment, infected cells were again washed three times in PBS and lysed with 1% Triton X-100 and 0.1% sodium dodecyl sulfate (SDS). Cell lysates were then serially diluted and plated on selective medium to assess bacterial counts.

Intracellular survival and replication in macrophages were carried out as previously described (25, 33, 35). Briefly, macrophages were grown in 24-well plates (approximately 5×10^5^ cells/well) for 24 hours. Bacteria were cultured in LB at 37°C to early stationary phase and the OD600 was adjusted to 0.1. The bacteria were opsonized in DMEM containing 10% normal mouse serum (Gemini Bio-Products, Woodland, CA) for 20 minutes at 37°C. Opsonized bacteria were used to infect J774 macrophages at an MOI of 10. Bacterial attachment and infection was enhanced by centrifugation at 500xg for 5 minutes. Infected macrophages were incubated further for 30 minutes at 37°C in 5% CO_2_. The macrophages were then washed twice with PBS to remove unbound bacteria and DMEM containing 10% FBS and 16 μg ml^-1^ gentamicin were added to the infection mixture. Two and twenty four hours after gentamicin treatment, infected macrophages were washed three times in PBS and lysed with 1% Triton X-100 and 0.1% SDS. Samples were then serially diluted and plated on selective media to enumerate the intracellular bacteria. The extent of replication was then determined by dividing the number of intracellular bacteria at twenty-four hour by the number at two hours. The data are the averages of three experiments performed in triplicate.

### Competitive infection in the mouse model

Wild-type and mutant *Salmonella* strains were grown overnight at 37°C in LB with proper aeration. Overnight bacterial culture was inoculated in fresh LB media containing 0.3M NaCl to an optical density of 1.0 (600 nm) at 37°C. At this point, the bacterial number was determined by serial dilution and plating on LB plates containing Streptomycin. Serial dilutions were then made in PBS to achieve the proper bacterial concentration for inoculation into the mice. Each mutant strain was mixed 1:1 with the wild-type SL1344 reference strain, and 200μL of the mixture were injected intraperitoneally into 6- to 8-week-old female BALB/c mice with the indicated inoculum. Two days after infection, mice were properly sacrificed, and their spleens and livers were harvested. Spleen and liver samples were homogenized and suspensions were serially diluted and plated on LB supplemented with streptomycin or streptomycin and kanamycin to enumerate bacterial numbers. The mutant strains are kanamycin resistant, which could grow on both plates; the wild-type *Salmonella* can only grow on LB with streptomycin. Mutant and wild-type organisms were enumerated and the competitive index (CI) was calculated as the percentage of mutant strain / wild-type *Salmonella* recovered as described previously (36, 37).

## Results

### Identification of *Salmonella* TTSS effector proteins

There are 5117 open reading frames in the genome of *Salmonella* SL1344 and 2357 of them (46%) are annotated as hypothetical or unknown functions. We reasoned that some of the hypothetical proteins might be *Salmonella* TTSS effectors. To identify novel *Salmonella* effectors, we chose to focus on proteins annotated as hypothetical of unknown function. A list of gene ontology search terms (**Table S2**) was compiled describing activities that could be or are currently known to be associated with *Salmonella* infections. In the bioinformatics analysis, functions of all proteins were first predicted using two independent algorithms, PFP (38) and ESG (39). Then, putative proteins that possess one or more of the search terms from the list were selected as candidates of effectors. This analysis led to the selection of 383 candidate TTSS effectors that fulfilled the selection criteria (**Table S3**).

### Identification of *Salmonella* effector proteins by the TEM1 reporter system

To determine whether any of the 383 candidate TTSS effectors are translocated into the host cell during *Salmonella* infection, we performed β-lactamase protein translocation assays on all 383 putative effector proteins. The N-terminus of each putative effector was translationally fused with the TEM1 protein using gateway clone technology. Briefly, entry vectors were obtained from Invitrogen carrying *Salmonella* putative effector genes flanked by AttL1 and AttL2 recombination sites. Destination vector was generated through inserting the recombination cassette (Invitrogen) into the plasmid carrying the gene encoding TEM1. The recombination cassette that contained the AttR1 and AttR2 sites was inserted directly upstream the gene encoding TEM1. Entry clone and destination vectors were transformed together into *E.coli.* Recombination between AttL (Entry) and AttR (Destination) sites lead to the transfer of the genes encoding the putative effectors into the destination vector resulting in the translational fusions of effector-TEM1.

To test whether these effector-TEM1 fusions were translocated during infection, HeLa cells were infected with effector-TEM1 fusion-expressing *Salmonella* strains for 15 min at an MOI of 10. Infected host cells were then loaded with CCF4-AM which excites at 409 nm and emits green fluorescence (520 nm). Translocation of the effector-TEM1 fusion proteins into the infected host cells leads to cleavage of the β-lactam ring of CCF4-AM. The cleaved product of CCF4-AM lead to the change in the fluorescence emission from green to blue (447 nm) when excited at 409nm. The translocation efficiencies were evaluated as percentage of green to blue conversions. *Salmonella* carrying plasmids expressing the SipA-TEM1 fusion was used as the positive control. Strains expressing effector-TEM1 fusion proteins that gave more than 5% of blue cells were retained for further analysis. Out of the 383 putative effectors, 22 were translocated into host cells (**Fig. 1**). To further confirm their translocation, the 22 effectors identified by the TEM1 assays were translationally fused with the CyaA tags (40–42). Similarly, infections were carried out with *Salmonella* carrying plasmids expressing the effector-CyaA fusions and the translocation efficiency was determined by measuring cellular cyclic AMP(cAMP) levels (40, 42). Out of the 22 putative effectors, 17 showed significant increase in cAMP levels when compared to the negative controls (**Fig. S1**). Further analysis indicated that STM3077 (YggG) is a putative Zn-dependent metalloprotease that contains the conserved HEXXH motif (**Fig. 2**).

**Fig. 1.**
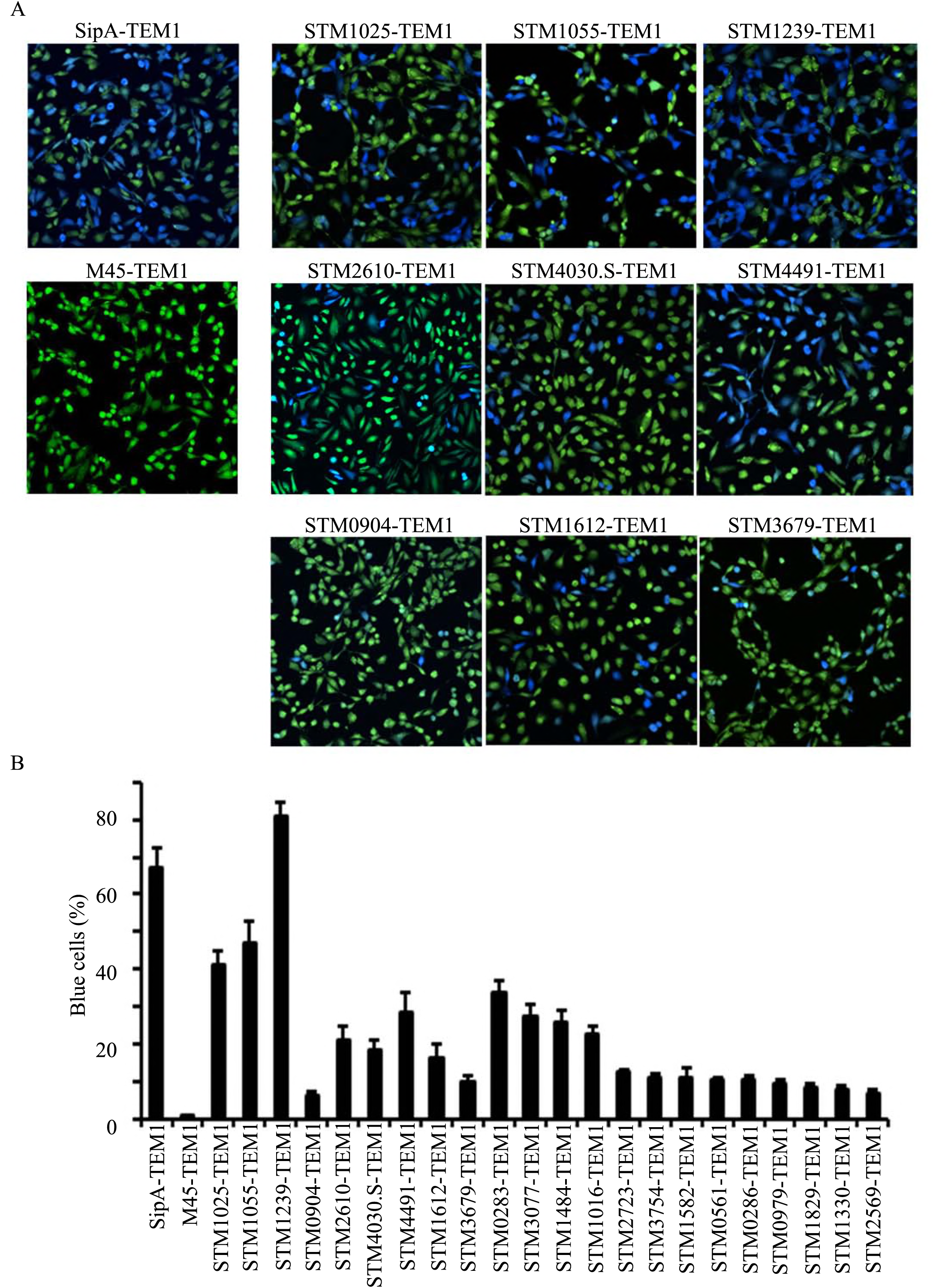
Translocation of *Salmonella* putative effector proteins into HeLa cells. (**A**) HeLa cells were infected for 15min with wild-type *Salmonella* carrying plasmid expressing the corresponding effector-TEM1 fusion proteins. Infected cells were then loaded with CCF4-AM, incubated at room temperature for 2h and then evaluated under a fluorescent microscope. Representative fluorescence micrographs were presented here showing the translocation of effector-TEM1 fusion proteins as indicated by the blue conversion from green. (**B**) Quantification of the translocation of *Salmonella* effector-TEM1 fusion proteins. Results shown are the mean ± SD of three independent experiments. M45-TEM1 was used as negative controls (green fluorescence) and SipA-TEM1 was used as a positive control.

**Fig. 2.**
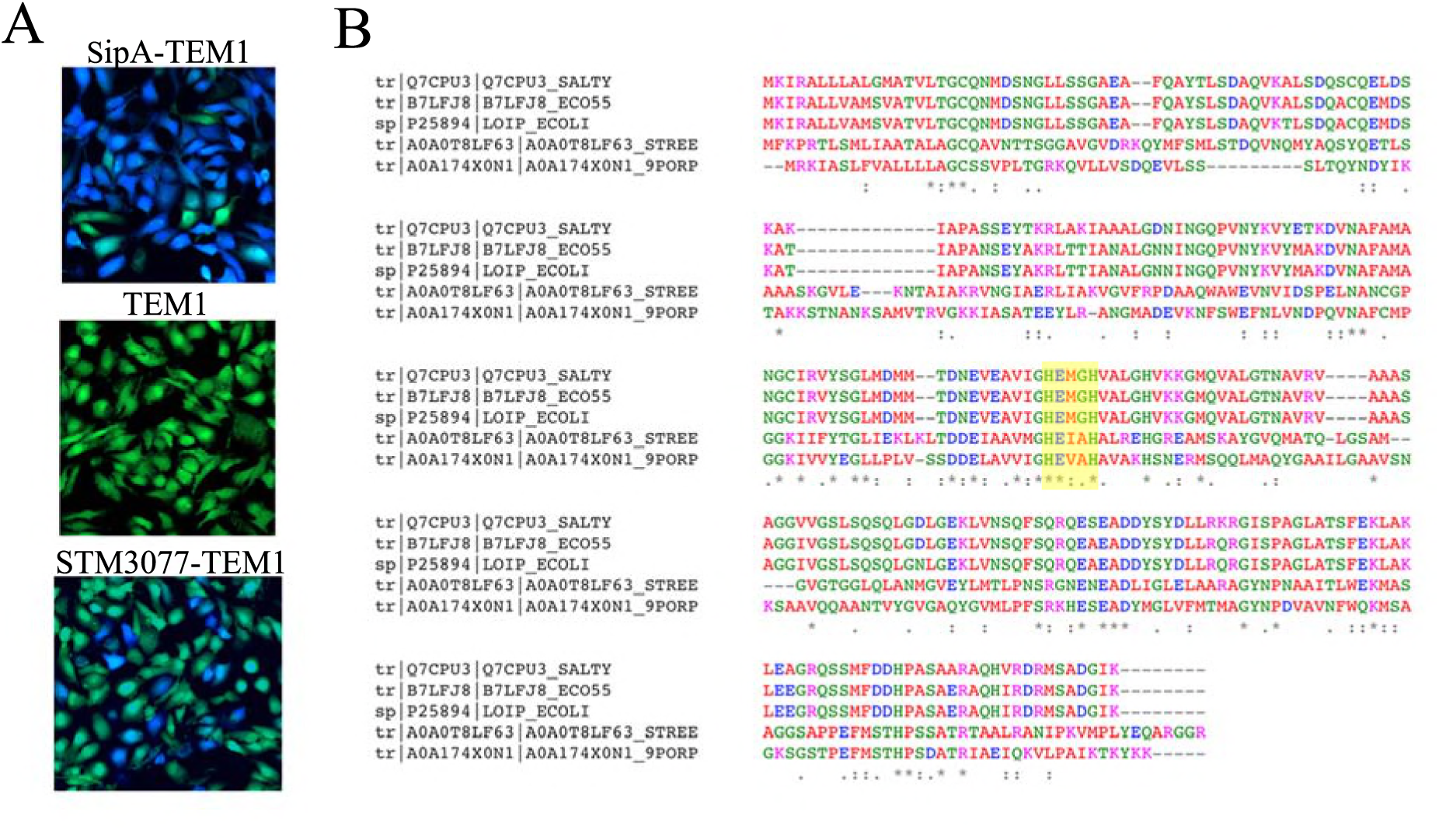
YggG contains active HEXXH domain and is predicted to be a metalloprotease. (**A**) Translocation of YggG into HeLa cells. HeLa cells were infected for 15min with wildtype *Salmonella* carrying plasmid expressing YggG-TEM1 fusion. Infected cells were processed and evaluated as described above. (**B**). Multi-sequence alignment of YggG with members of M48 metalloprotease family. Top to bottom: YggG (*Salmonella* Typhimurium), YggG (*Escherichia coli*), Metalloprotease LoiP (*Escherichia coli*), Uncharacterized metalloprotease YggG (*Streptococcus pneumoniae*), Uncharacterized metalloprotease YggG (*Parabacteroides distasonis*). The HEXXH motif is highlighted in yellow.

### YggG is cytotoxic to mammalian cells

*Salmonella* effectors are translocated into the infected host cells where they exert their effects by modulating the host cell function, such as the actin cytoskeleton rearrangements, and the inflammatory responses. The pathogens usually maintain a balance of these functions to avoid detrimental effects, such as cell death, during infections. However, when these effectors are artificially overexpressed, some of them are cytotoxic to the mammalian cells. Thus, we overexpressed the 22 effectors in mammalian cells and examined them for toxicity. One of the newly identified effector YggG was toxic to mammalian cell upon overexpression. Cells transfected with GFP-YggG detach from the plate surface and round up (**Fig. 3A**), typical of cytotoxicity. To quantitatively measure the cytotoxicity induced by GFP-YggG overexpression, the lactate dehydrogenase (LDH) release assay was performed according to manufacturer’s instruction. Overexpression of the wild type GFP-YggG resulted in an increase in LDH release 24 hours post transfection (**Fig. 3B**). To test whether the putative Zn-dependent metalloprotease is required for the observed cytotoxicity, a GFP-YggG^E131C^ mutant (point mutation in the conserved HEXXH motif) was constructed and similarly overexpressed as described above. The overexpression of the GFP-YggG^E131C^ resulted in much reduced cytotoxicity comparing to that induced by the wild type GFP-YggG (**Fig. 3B & 3C**). These data further validated YggG’s cytotoxicity to mammalian cells and its protease activity is responsible for the cytotoxic phenotype.

**Fig. 3.**
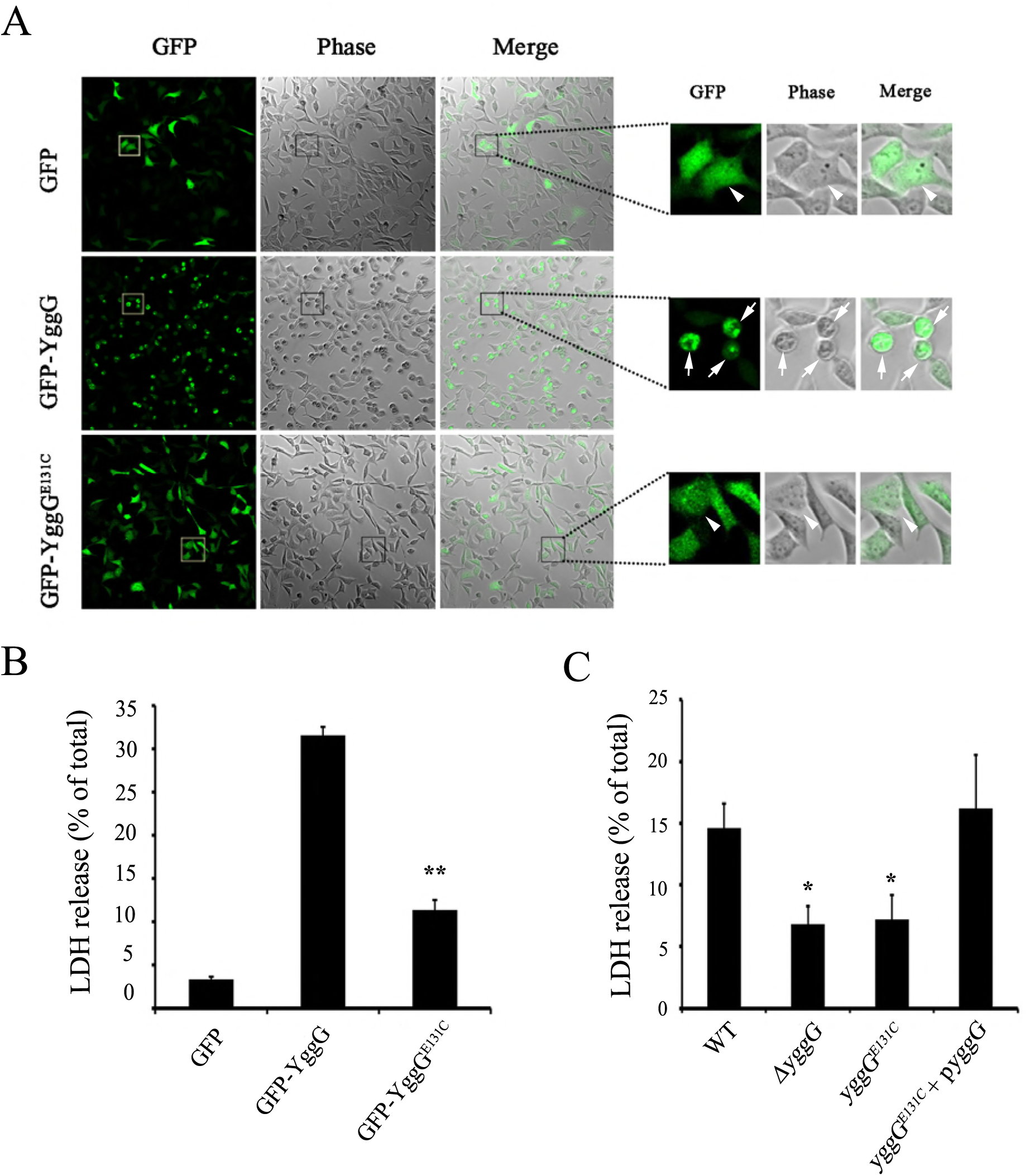
YggG is toxic to mammalian cells. (**A**) Plasmids expressing GFP, GFP-YggG or GFP-YggG^E131C^ fusion proteins were transfected into 293A cells. Cell morphology was then analyzed by confocal diffraction phase microscopy. Arrowheads indicate normally shaped cells, while arrows indicate cells that are rounded-up (signs of toxicity). (**B**) Plasmids expressing GFP, GFP-YggG or GFP-YggG^E131C^ were transfected into 293A cells for 24 hrs. Cell cytotoxicity was quantified using LDH-release assay as described in the methods. (**C**) HeLa cells were infected with wild-type *Salmonella*, *yggG* null mutant strain (Δ*yggG*), chromosomal point mutant strain (*yggG^E131C^*), or the *yggG^E131C^* strain carrying a plasmid expressing the wild-type YggG (*yggG^E131C^*+p*yggG*). Cell cytotoxicity was quantified using the LDH-release assay. Error bars indicate standard deviations of triplicate experiments.

### YggG does not affect *Salmonella* invasion and replication *in vitro*

Entry of *Salmonella* into epithelial cells is an important virulence trait and one of the first steps to initiate *Salmonella* infection. With the help of various effector proteins, *Salmonella* actively manipulates host cell actin dynamics to promote the engulfment of the bacterium. To determine if YggG is necessary for bacterial invasion, HeLa cells were infected with the wild type or *yggG* mutant strain for 30 min at 37°C, and the *Salmonella* invasion efficiency was analyzed using an inside-outside staining assay. The *invA* (deficient in SPI-1 function) strain was used as a negative control. The invasion efficiency of the *yggG* mutant strain was similar to that of the wild-type strain (**Fig. 4A**). This result suggests that YggG is not necessary for *Salmonella* invasion. Following entry into infected cells, the bacteria must survive and replicate within macrophages in order to cause systematic infections. To test the ability of the *yggG* mutant strain to survive and replicate within macrophages, a gentamicin protection assay was performed using macrophage cell line J774. The *ssaV* (deficient in SPI-2 secretion) mutant strain was used as a negative control. As shown in **Fig. 4B**, the *yggG* mutant strain replicated to a similar degree as that of the wild-type strain, suggesting that YggG does not directly affect the survival/replication of *Salmonella* in J774 macrophages.

**Fig. 4.**
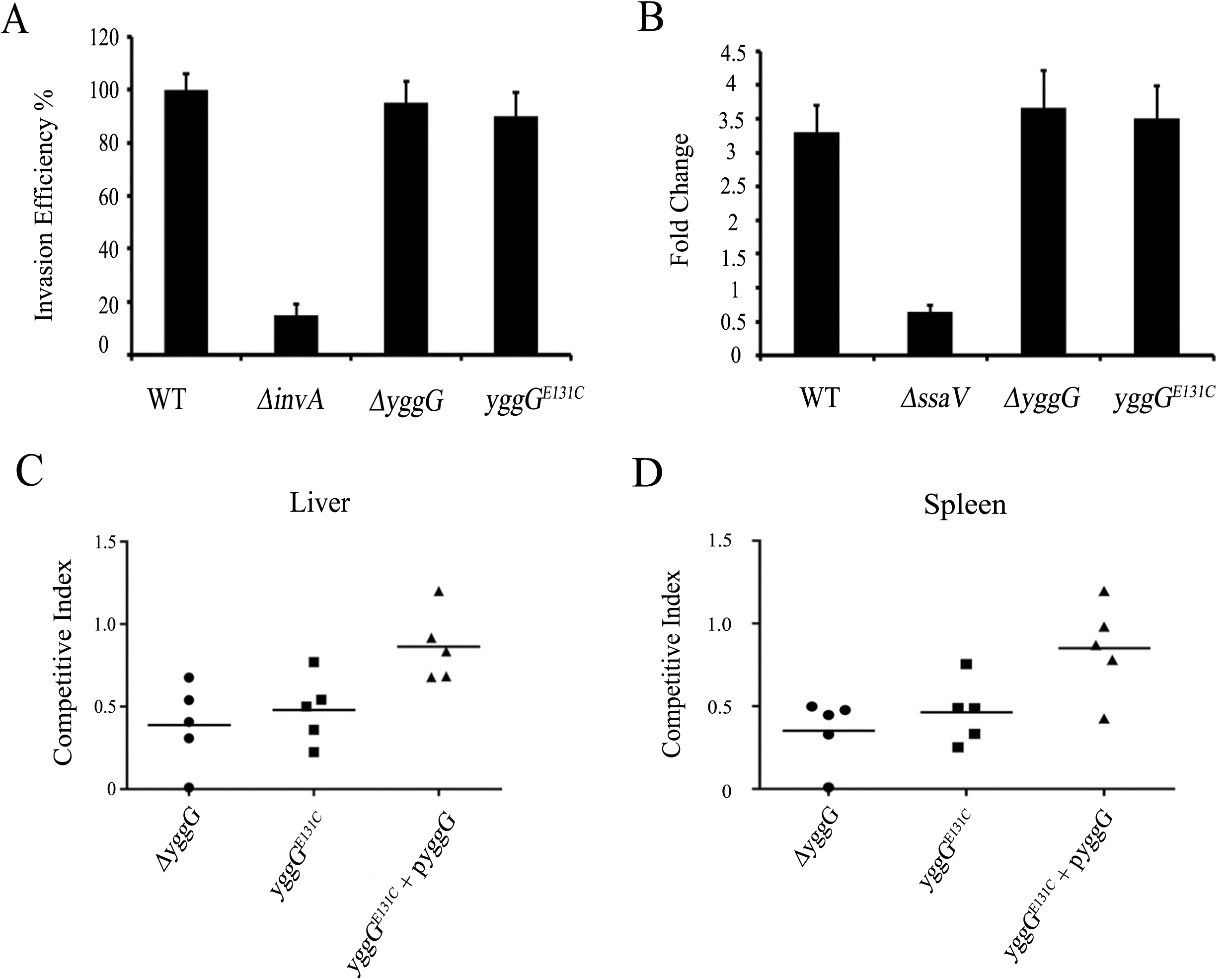
YggG is essential for *Salmonella* virulence in mice. **(A)** HeLa cells were infected with wild-type *Salmonella*, or indicated mutant strains at an MOI of 10 for 15 minutes. Invasion efficiency of the wild-type *Salmonella* was artificially set to 100% and the other strains were compared to that of the wild-type strain. A invasion deficient *invA* mutant was used as the negative control. Error bars indicate standard deviations of triplicate experiments. **(B)** J774 macrophages were infected with indicated strains and the replication efficiency was evaluated by bacterial count as described in the methods. The SPI-2 deficient mutant strain (*ssaV*) was used as the negative control for replication in the macrophages. Error bars indicate standard deviations of triplicate experiments. **(C & D)** BALB/c mice were inoculated intraperitoneally with a 1:1 mixture of wild-type and indicated mutant strains. Liver (C) and spleen (D) were collected 48 hrs post-inoculation and bacteria loads were enumerated. The C.I. were calculated as described in the methods. Each symbol represents a mouse, and horizontal bars correspond to the means. A one-sample t-test was used to determine whether a value was significantly different of one.

### YggG contributes to *Salmonella* virulence in mice

*Salmonella* Typhimurium causes typhoid-like disease in mice, the mouse model of infection has been used to study acute *Salmonella* infections. Despite the fact that typhoid-like symptoms in the mice are different from the gastrointestinal inflammation in humans, the mouse model has been widely used to characterize virulence genes relevant for bacterial invasion, survival and replication that are essential for causing diseases in humans (3, 6). To test whether YggG plays a role in *Salmonella* survival and replication, we employed the well-documented competitive index (CI) assay using the mouse model of infection (36, 37). Wild type *Salmonella* or the *yggG* (Δ*yggG*) mutant strain carrying the kanamycin resistant gene were mixed in 1:1 ratio before infecting the mice. Groups of five BALB/c mice were intraperitoneally infected with approximately 10^4^ CFU of the 1:1 mixture of the wild type and Δ*yggG* strains. Mice were euthanized two days post infection to enumerate bacterial numbers in both the livers and spleens. The CI values were then calculated as described previously (36, 37). Our data showed that the CI of Δ*yggG* is significantly lower than 1 (**Fig. 4C & 4D**), indicating that YggG is essential for *Salmonella* virulence in the mouse model of infection. To further test if the putative Zn-dependent metalloprotease of YggG is required for the virulence in mice, we first constructed a strain *yggG^E131C^*, with a chromosomal point mutation in the conserved HEXXH motif, carrying the kanamycin resistant gene. When tested using the competitive index assay described above, the *yggG^E131C^* strain is attenuated in virulence similar to that of the Δ*yggG* null mutant strain. To further make certain that the attenuated virulence observed was indeed due to YggG, complementation experiment was carried out by introducing a plasmid harboring wild type *yggG* into the *yggG^E131C^* strain (*yggG^E131C^*+p*yggG*). Our data showed that the introduction of the wild type *yggG* restored the virulence defect of the *yggG^E131C^* strain (**Fig. 4C & 4D**).

## Discussion

*Salmonella* ensures its colonization and proliferation in the host by secreting an array of effector molecules that modulate various cellular functions. The identification and characterization of these effector proteins are crucial to advance our understanding of *Salmonella* pathogenesis. Despite the progress in identifying a number of key effectors and in dissecting their biochemical and cellular activities, the molecular mechanisms of how *Salmonella* modulate host cell trafficking and facilitating subsequent replication remain largely unknown (43–45). Although many of these functions are known to be conferred by the *Salmonella* TTSSs, effectors that are responsible remain elusive. For example, the delayed macrophage cytotoxicity (21) and the avoidance of oxidative burst (17, 22–24) during *Salmonella* infection has not been understood.

It is known that type III secreted effectors do not share sequence similarity and are difficult to predict based on their amino acid sequences (30). Many experimental approaches have been developed to identify secreted effectors in *Salmonella.* Geddes *et al.* (2005) introduced the mini-Tn5-cycler to the *Salmonella* chromosome to generate translational fusions to the CyaA reporter enzyme. Putative effectors were identified based on cAMP levels in infected cells (17). They successfully found three new effectors and seven that have been previously reported. Choy *et al* (2004) identified novel *Salmonella* effectors by comparing *Salmonella* proteins to known secreted proteins from enterohemorrhagic *Escherichia coli* and *Citrobacter rodentium* (46). Niemann et al performed a proteomic analysis and identified effector proteins secreted into defined minimal medium under SPI-2 TTSS conditions (47). As a result, they identified 12 candidate SPI-2 effector proteins many of which were shown to be required for full *Salmonella* virulence in mice. Six mutants were significantly attenuated for spleen colonization. In addition, Cheng et al have recently identified a novel SPI-1 effector, SopF that is required for *Salmonella* virulence in mice using a similar proteomic approach (48).

In this study, we performed a large-scale screening of *Salmonella* novel effectors with a focus on unknown function proteins in the *Salmonella* genome. We coupled the power of the bioinformatics predictions with experimental protein translocation reporter assays to identify novel *Salmonella* effectors. Among the 383 putative effector proteins screened, we identified 22 novel effectors that are translocated into host cells during *Salmonella* infection using a β-lactamase protein translocation reporter assay. Interestingly, one of them was reported to be a type III effector by another group (17). Among the newly identified effectors, one of the effector YggG induces host cell death upon overexpression, suggesting YggG affiliated with properties that are responsible for the cytotoxicity. Further experiment showed that the protease activity of YggG is responsible for the cytotoxic phenotype. Moreover, the protease activity of YggG was found to be essential for *Salmonella* virulence in the mouse infection model.

Many proteases are known to be involved in *Salmonella* virulence (49–55). The lon protease has been shown to negatively regulate the efficiency of invasion of epithelial cells and the mutant was attenuated when administered either orally or intraperitoneally to mice. It was also shown that the lon-deficient mutant strain failed to survive and grow in macrophages (49, 54, 55). The ClpXP protease was proposed to suppress macrophage apoptosis by regulating the expression of SPI-1 and flagella (56). Moreover, host proteases, such as caspases are also known to play important roles during *Salmonella* infections (51, 53, 57).

Although cell culture models have been widely used to study *Salmonella* invasion and survival, animal infection models are often irreplaceable due to their intricate immune systems (58–62). Our study has demonstrated that YggG and its protease activity are required for *Salmonella* virulence in the mouse infection model. Surprisingly, no detectable roles in invasion into epithelial cells and no effect on the survival and replication inside cultured macrophages. *Salmonella* infection involves multiple processes in which the bacteria encounter and compete in order to survive and replicate. In addition, *Salmonella* has to gain a growth advantage over the intestinal microflora while inducing inflammation in the gut (63–66). Our data does not provide the precise molecular mechanism on how YggG function to promote *Salmonella* virulence. It is tempting to speculate that YggG may be involved in altering some aspects of the host immune system to promote infection. The precise molecular and cellular mechanism require further investigation.

## Figure Legends

**Fig. S1.** Translocation of *Salmonella* putative effector proteins fused with CyaA. *S*. Typhimurium strain SL1344 expressing the CyaA fusions were used to infect HeLa cells. The cAMP levels of infected cells was determined. SipA-CyaA fusion was used as positive control while vector expressing CyaA alone was used as a negative control.

